# Cuticulome.org: a centralized database of proteins in arthropod cuticles

**DOI:** 10.64898/2026.10.18.759974

**Authors:** Alex Wardale, Naomi Nakayama, Cédric Finet

## Abstract

Cuticle proteins, especially the cuticular protein (CP) multigene family, are essential for the building, maintenance, and environmental adaptation of the arthropod exoskeleton. The extensive size of CP repertoire, which often represents 1-2% of the protein-coding genes, is a challenge. The discontinuation of previous centralized databases for CPs and the lack of consistent nomenclature in the field have hindered functional analyses and comparative studies. Here, we present Cuticulome.org (accessible at https://www.Cuticulome.org), a curated knowledgebase of proteins that are found in arthropod cuticles. Database entries include protein sequences, tissue and developmental stage specificity, and diverse functional annotations where available, while implementing a standardized nomenclature system to facilitate intra- and cross-species analyses. At Cuticulome.org, users can download complete taxon-specific cuticular proteomes, entire CP families, or build their own custom datasets defined by taxonomic, protein family, functional, or keyword-based criteria. Cuticulome.org also offers two integrated sequence analysis tools: *miniBLAST*, which allows users to identify putative orthologous CPs, and *Cuticular Protein Classifier*, which accurately classifies a protein sequence to an established CP family based on custom Hidden Markov Models. Together, these features establish Cuticulome.org as the new centralized resource for systematic analysis of cuticle proteins.

## INTRODUCTION

Arthropods are covered with a non-living cuticle, or exoskeleton, which is secreted by a single-layered epidermis. The cuticle is a composite material consisting of chitin fibres, proteins, pigments, lipids, and occasionally metal ions and mineral crystals (Muthukrishnan *et al*. 2022). Although early efforts have revealed the hierarchical organization of chitin into nanofibrils and thicker fibres, the precise structural significance of the remaining components has only recently been identified. Pigments, for example, have been proven to alter insect coloration in tandem with morphology (Matsuoka and Monteiro 2018)(Banerjee *et al*. 2024)(Chatterjee *et al*. 2026) and mechanical properties (Massey *et al*. 2019)(Balakrishnan *et al*. 2025). Metal ions and mineral crystals have been repeatedly used to strengthen cuticle by acting, respectively, as cross-linking agents for proteins and chitin coordinate (Politi *et al*. 2012)(Sakr *et al*. 2025) and through biomineralization (Rebora *et al*. 2023)(Vittori 2024). However, proteins identified in arthropod cuticles are arguably the most promising class for new discoveries in biology and material bioengineering (Willis *et al*. 2012)(Miserez, Yu, and Mohammadi 2023), with some examples of insect cuticle-inspired hydrogels and bioplastics (Chen *et al*. 2024)(Wu *et al*. 2026).

Even though the cuticulome contains a variety of proteins, including enzymes and lipoproteins, cuticular proteins (CPs) remain the predominant class with CP genes accounting for >1-2% of the total insect genome (Cornman *et al*. 2008)(Willis 2010). CPs are a diverse family of structural proteins secreted by epidermal cells that combine with chitin to form the multifunctional arthropod cuticle (Rebers and Willis 2001)(Hu *et al*. 2025). They are essential for exoskeleton development, environmental adaptation, and immunity (Charles 2010)(Mallick and Eleftherianos 2024)(Qi and Liu 2025). Andersen and colleagues’ groundbreaking work on CP extraction and sequencing paved the way for CP research (Andersen, Hojrup, and Roepstorff 1995), which was followed by Willis’ work on CP nomenclature and annotation (Willis 2010). Since then, our understanding of CPs has been hampered by methodological and experimental difficulties due to their large number, the presence of unpredictable intrinsically disordered regions and low-complexity amino acid sequences, and functional redundancy. However, with the advent of next-generation sequencing and universal genome editing technologies (e.g., CRISPR-Cas9), numerous studies on the identification and functional characterization of CPs have been published during the last decade. The rapid growth in our understanding of CP functions is difficult for current topical studies to comprehensively address (Qi and Liu 2025). This is the rationale for creating a database.

A first relational database of arthropod CPs, cuticleDB, was created twenty years ago with the goal of identifying CPs in a few available genomes (*Anopheles gambiae*, *Apis mellifera*, *Bombyx mori*, *Drosophila melanogaster*, *Nasonia vitripennis*), as well as providing an online tool for predicting Rebers and Riddiford (R&R) consensus RR-1 and RR-2 domains (Magkrioti *et al*. 2004). However, this database was last updated in 2009, and the amount of genomic data has grown tremendously since then. In parallel, the online tool CutProtFam-Pred was developed to identify and classify putative CPs relying on legacy 2014 profile Hidden Markov Models (HMMs) (Ioannidou *et al*. 2014). While highly effective at their launch, these models cannot accurately account for the massive explosion of recently discovered arthropod genomes and the identification of completely novel CP families.

Our new database, Cuticulome.org, fulfils several functions: (1) compiling the functional data on natural and engineered CPs, (2) creating datasets of comprehensive CP repertoire for some species, (3) offering tools such as *miniBLAST* and *Cuticular Protein Classifier* that broadens the strict identification of RR-1 and RR-2 motifs, and (4) enabling community contributions through a submission portal. Our database aims at standardizing nomenclature of CPs and facilitating comparative studies.

## MATERIALS AND METHODS

### Literature screening and data curation

Full taxonomic data (class, species) were extracted from NCBI Taxonomy (Schoch *et al*. 2020). Expression and functional data were collected by screening over 200 CP research papers published between 1986 and 2026. From these, we extracted data relating to the expression in space (organs and/or tissues) and time (embryo, larva, pupa/nymph, adult), as well as functional information, where available. Proteins with experimentally validated functions were classified as function-defined, whereas proteins without this evidence were kept as non-function-defined entries. Protein and coding sequence data were also collected from the NCBI accession numbers that were cited in the publications. When accession numbers were not mentioned or were outdated, sequences were obtained by further investigating the associated genome or transcriptome resources, dedicated databases of the specific species (such as FlyBase for *Drosophila* species, and SilkBase for *Bombyx mori*), or by contacting the authors directly. Species-level cuticular proteomes were curated from studies that implemented transcriptome- or proteome-wide approaches to identify larger sets of CPs. We reviewed and included repertoires of diverse orders to maximize taxonomic breadth across arthropods, and when multiple papers were available for the same species, we cross-checked entries and included those with the most optimal annotation (based on recency and sequence completeness). In addition to natural CPs, we added to our database engineered ones and their respective sequences (Zheng *et al*. 2025).

### Nomenclature standardization

The lack of a standardized naming system in the field has led to inconsistent nomenclature when identifying homologous CPs between arthropod species. The same protein can present different formats across different species, such as *Dmel_CPR1*, *Dscpr1*, and *MpCPR1* for CPR1 in *Drosophila melanogaster*, *Drosophila suzukii*, and *Myzus persicae*, respectively. Additionally, some CPs have been reported independently with different names and only later recognized as the same protein, resulting in duplicate nomenclature such as BmPCP and BmorCPH31 in *B. mori*. This inconsistency is problematic not only for comparative analyses, but also for database searches and cross-species functional studies. To address this, Cuticulome.org implements a new simplified nomenclature system for CP entries, while also preserving searchability for previous naming systems, alternative names, and synonyms. The format follows a three-letter species code followed by an underscore separator and the shortened protein name: *Xxx_ProteinName*. The species code consists of the first letter of the genus (capitalized) followed by the first and second letter of the species (both lower case). For example, following this system, CPR1 in *D. melanogaster* would be formatted as *Dme_CPR1* in the database entry.

Seldom, naming can overlap, as would be the case for *Heliothis virescens* and *Heortia vitessoides*, CPR1 of which would identically be *Hvi_CPR1* following our standardized nomenclature system. To avoid confusion in these cases, the entire genus name is included as a prefix in the database entry and export to avoid any ambiguity in downstream analyses. Following this, CPR1 would be identified as *Heliothis_Hvi_CPR1* and *Heortia_Hvi_CPR1* for these two species, respectively, with both options being filtered in the case of searching for the three-letter prefix.

We note that this naming standardization applies solely within Cuticulome.org for simplified searching and is not intended to replace any official nomenclature.

#### Cuticular Protein Classifier development

We first curated and grouped protein sequences for each established CP family and used them as seed datasets to generate family-specific multiple sequence alignments. These alignments were then used to build custom Cuticulome.org profile HMMs using the open-source HMMER3 software package (Eddy 2011). The resulting HMMs represented conserved CP family sequence features, while still considering the known motif structures, domain organization, and in some cases, taxonomic limitations. For each CP family HMM, stringent E-value and score thresholds were set to reduce false-positive classifications while retaining sensitivity for validated and classified CPs. In several cases, proteins that were not included in the original seed datasets were later recovered by the HMM as potential candidates and were subsequently confirmed as bona fide members of the CP family they were classified into after manual inspection and literature verification. One example of this was *Tenebrio molitor* LPCP-22, which was not initially included in the CPF seed dataset as it was collected from separate literature that had identified it as a low complexity CP (Rondot 1995). Tmo_LPCP-22 was later recovered by our CPF HMM as an unclassified potential candidate. During classification verification, we identified that not only did it match the diagnostic CPF sequence feature, but it was also described in another study, including the conserved internal CPF region and CPF-like C-terminal (Togawa *et al*. 2007), further supporting the validity of our classifications.

An archival copy of the code, curated database seed files, reasoning, and associated resources and metadata used for the profile HMM-based models for *Cuticular Protein Classifier* have been deposited and are available in Zenodo (https://doi.org/10.5281/zenodo.20150640).

### Backend implementation

Cuticulome.org is built around a structured SQLite database of curated literature-derived CP data and sequence information. The database is maintained through the dedicated Cuticulome.org GitHub repository (https://github.com/Wardale24/Cuticulome.org) and serves as the source for both entry retrieval and the two integrated sequence-analysis tools. These tools are both provided through a dedicated backend which is hosted on Render.com.

The *miniBLAST* tool backend implements BLASTP from the NCBI BLAST+ suite (Camacho *et al*. 2009). Queried protein sequences are searched against the curated Cuticulome.org protein sequence dataset using an E-value threshold of 1×10 ³, which was selected to balance sensitivity for less conserved CPs with the exclusion of weak matches expected to occur frequently by chance. The results are ranked by sequence matches and presented with associated entry information.

The *Cuticular Protein Classifier* uses the aforementioned custom Cuticulome.org profile Hidden Markov Models developed for established CP families. Queried amino acid sequences are compared against the models, and the classifier outputs the detected HMM evidence. This evidence is presented as the predicted CP family with supporting HMM matches, an overall confidence level, and its associated statistics.

### Frontend implementation

The Cuticulome.org knowledgebase is accessible via a dedicated web interface that is continuously developed with a user-centered perspective. We designed the interface to emphasize searchability, flexible filtering, and the generation of customizable and downloadable datasets for downstream analyses. In this way, users can retrieve CP-related datasets according to different research interests, whether it is species-level repertoires, protein-family datasets, function-associated entries, or customized datasets generated from selected filters. To improve webpage navigation and usability, we performed extensive bug testing with external web developers. This testing was primarily used to identify issues related to layout, navigation, and general user experience, while attempting to identify any type of error or accessibility issue. We also inquired members of the scientific community regarding the ease of use of Cuticulome.org, implementing their feedback to further improve the web interface.

## RESULTS

### Database content and coverage

To date, Cuticulome.org contains 2678 unique CP records, of which 349 have experimentally validated functions in arthropods. Each CP record represents a standardized protein entry, structure of which is summarized in **Table 1**, which includes database identifiers, alternative naming, taxonomic information, protein-family, NCBI protein sequence accession numbers, curated expression and functional annotations, and literature references. The database covers 35 protein families across 66 species, 53 genera, and 14 orders, providing a centralized resource for exploring both function-defined and non-function-defined CPs.

**Table 1.** Curated metadata fields included in Cuticulome.org protein entries.

| Data type |
| --- |
| Standardized cuticular protein name |
| Alternative naming |
| Taxonomic information |
| Cuticular protein family |
| Expression |
| Tissue |
| Timing |
| Variation under treatments |
| Details |
| Molecular function |
| Biological process |
| Biological function |
| Molecular mechanism |
| Source reference |
| NCBI/Genbank entries |

**Table 2.** Top 8 species represented in Cuticulome.org by number of CP entries. Only one Drosophila species is included to better demonstrate the diversity of species represented in Cuticulome.org and avoid overrepresentation of a single genus.

| Species | Number of CP entries |
| --- | --- |
| *Bombyx mori* | 300 |
| *Anopheles gambiae* | 285 |
| *Daphnia magna* | 262 |
| *Drosophila melanogaster* | 168 |
| *Gerris buenoi* | 155 |
| *Nilaparvata lugens* | 141 |
| *Cryptolestes ferrugineus* | 131 |
| *Frankliniella occidentalis* | 128 |

Taxonomic representation in Cuticulome.org reflects the distribution of currently available arthropod CP studies, which is skewed toward Insecta (**Figure 1A**), with most species represented in the database belonging to this class (89.4%), while a smaller portion is described in Arachnida (7.6%) and other arthropod classes (3%). In terms of unique CP records, the most represented species is *B. mori* with 300 proteins, closely followed by *Anopheles gambiae* (285 CPs), *Daphnia magna* (262 CPs), and *D. melanogaster* (168 CPs) (**Figure 1B**).

**Figure 1.**
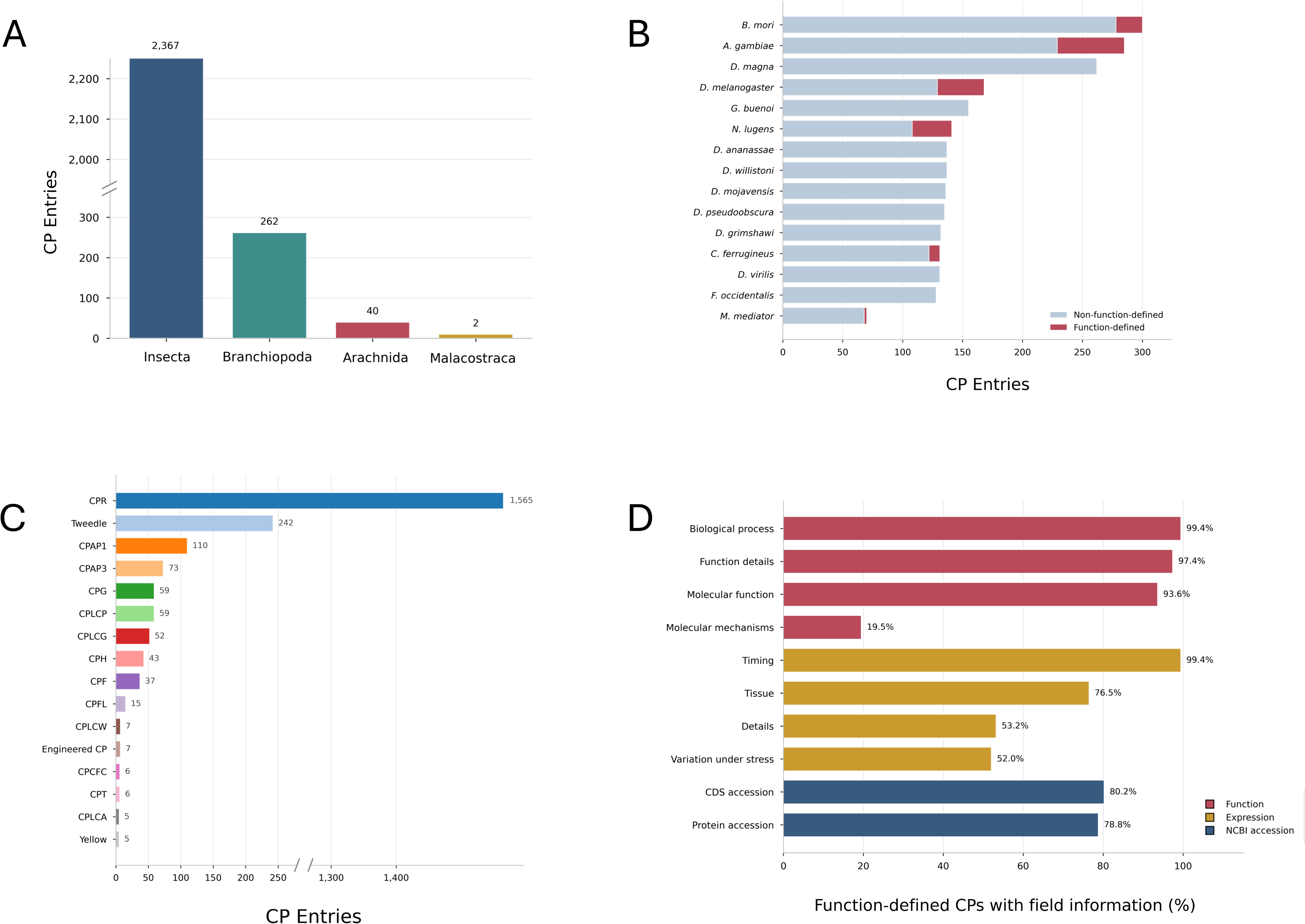
Cuticulome.org content and coverage overview. (A) Distribution of Cuticulome.org CP entries across represented arthropod classes. (B) Species with the largest number of curated CP entries, separated into function defined and non-function-defined records. (C) Distribution of CP entries across established CP families. CPR subclasses (RR-1, RR-2, RR-3) were grouped under CPR for visualization. (D) Completeness of curated annotation fields for function-defined CPs, showing the proportion of entries with curated functional, expression, and accession information.

In terms of established CP families, the largest number of CPs included in Cuticulome.org belongs to CPR, with 1560 proteins (**Figure 1C**), of which 756 have been subclassified into CPR RR-1 and CPR RR-2 protein families. The next most represented CP families are Tweedle (242 CPs), and CPAP (188 CPs). Synthetic CPs were classified as “Engineered CP”.

For function-defined CPs, annotation completeness was further assessed across selected functional, expression-related, and accession fields, highlighting the extent to which experimentally characterized proteins are supported by curated biological descriptions, expression information, molecular mechanism details, and NCBI sequence identifiers (**Figure 1D**).

### Database browsing and customizable dataset downloads

To maximize accessibility for a wide range of users, we have made the curated CP data searchable through three complementary browsing routes which allow for both targeted searches and broader comparative analyses: by CP, by protein family, and by species. The CP browsing page (https://www.Cuticulome.org/browse) allows users to search for proteins based on Cuticulome.org standardized nomenclature, original protein name, NCBI accession number, and functional keywords. Each CP entry links to a dedicated record page containing the available associated information, including amino acid sequence, curated expression and functional annotations, alternative names, and literature references.

In addition to CP-level browsing, Cuticulome.org includes family- and species-centered views. Protein family pages summarize, in unique boxed entries, the number of CPs, species, and average sequence length pertaining to each established CP family, while also providing highlighted examples of CPs with validated function where available (https://www.Cuticulome.org/families). Species pages on the other hand, provide access to the entire cuticular proteome that is available on Cuticulome.org for each species, allowing users to easily filter their organism of interest and obtain summaries of their corresponding CP records (https://www.Cuticulome.org/species). These different browsing routes ultimately allow users to easily navigate individual CP entries, established protein family datasets, and species-level repertoires without requiring troublesome webpage interaction or database filtering.

The browsing system in Cuticulome.org is directly linked to dataset export. Through the Download page, users can build custom datasets by filtering CP entries according to genus, species, protein family, functional annotation status, keyword-based search criteria, or a mixture of these (https://www.Cuticulome.org/downloads). Once filtered, datasets can then be exported in two complementary formats: a single FASTA file containing all amino acid sequences for entries filtered with a standardized and provenance-informative header, and a CSV file containing the associated curated metadata, expression and functional annotations, NCBI accession numbers, and literature references. This allows users to access tailored datasets that directly address their needs and download FASTA files that can be directly incorporated into downstream analyses without additional formatting.

### Integrated sequence analysis tools

To further support CP research, Cuticulome.org offers two integrated sequence analysis and classification tools without the need for local installation of alignment software or any familiarity with the command line.

#### miniBLAST

*miniBLAST* is a lightweight sequence-similarity search tool that allows users to query the Cuticulome.org database with a protein sequence of interest to identify identical, homologous, or related CPs across arthropod species (https://Cuticulome.org/tools/miniblast). *miniBLAST* accepts amino acid sequences in raw or FASTA format and searches them against the curated set of both function-defined and non-function-defined CPs in Cuticulome.org.

*miniBLAST* outputs the Cuticulome.org standardized protein name, species, NCBI accession number, percentage sequence identity, query coverage, E-value, and bit score for each returned hit. In addition, a functional-status indicator is displayed next to each result, allowing users to discern between matches that have an experimentally validated function from those that do not. Users can additionally download the results as a spreadsheet which includes all associated database information for each matching entry.

#### Cuticular Protein Classifier

*Cuticular Protein Classifier* supports family-level annotation of query protein sequences (https://Cuticulome.org/tools/classifier). This tool allows users to submit an amino acid sequence and evaluate whether it matches sequence signatures associated with established CP families, which are calculated and assigned based on custom Cuticulome.org profile HMMs.

After querying an amino acid sequence, the classifier returns the predicted CP family, an overall confidence level, and the supporting HMM evidence used to generate the classification, including the detected model matches and its associated statistics. The tool is particularly useful for evaluating newly identified or poorly annotated CPs. However, as CP classification may depend on motif structure, domain organization, sequence completeness, and in some cases broader biological or taxonomical context (such as CPLCW, only identified in mosquitoes at the time of this publication), classifier results should be interpreted as supporting evidence rather than definitive annotations.

### User-contributed annotations and datasets

We have thoroughly screened publications to include a variety of functional and expression-related annotations on CPs, with the aim of maximizing the breadth and depth of curated functional information. In addition, we continue to add CPs by sifting through database entries of different arthropod species and keeping up with the growing number of recent CP-related publications. However, curating the database remains an ongoing effort that requires community collaboration and input. To facilitate this, Cuticulome.org welcomes contributions by including a user submission page that allows community members to add newly characterized arthropod CPs to the database (https://Cuticulome.org/submit). This page includes a submission form with fields that refer to protein name, species, accession number, sequence, publication details, and several functional- and expression-related annotations. All submitted entries are reviewed by the Cuticulome.org curators before entering them into the database. During this process, the submitted proteins are evaluated for validity and are assigned a name based on the standardized nomenclature of Cuticulome.org. For larger batch submissions, users can access the Contact us page (https://Cuticulome.org/contact) where they can directly communicate with curators and avoid repetitive manual submissions. Through Cuticulome.org we aim to support the CP field by enabling researchers to easily access information and datasets of their choosing to advance in their investigations. This includes supporting the annotation and classification of putative CPs, enhancing the database, and collaboration opportunities.

### USE-CASES

Below, we describe two use-case examples that demonstrate how Cuticulome.org can support CP investigation, annotation, and comparative analyses. These workflows are not only intended for researchers that are already working with CPs, but also for members of the broader research community who could possibly retrieve cuticle proteins from omic datasets. The first use-case illustrates how a user can evaluate whether a newly identified protein sequence belongs to an established CP family and identify related proteins that are already present in the database. The second exemplifies how Cuticulome.org can be used to quickly retrieve customized CP datasets to directly plug into downstream comparative or phylogenetic analyses.

### Example 1: characterization of a newly identified CP candidate

Both transcriptomic and proteomic studies frequently recover protein sequences that are poorly annotated or only broadly described as cuticle-associated proteins. In some cases, researchers may not know whether an amino acid sequence is a CP at all, and even if it were, the next posed questions could be if it belongs to an established CP family, if there are similar proteins described in other species, or if similar CPs have related validated functions.

As an example, a user can submit an amino acid sequence of a candidate protein to both *miniBLAST* (**Figure 2A**) and *Cuticular Protein Classifier* (**Figure 2B**). The former returns ranked matches with the Cuticulome.org standardized nomenclature (**Figure 2C**), whether the matched protein is function-defined, and the associated match statistics. The latter returns a predicted CP family based on Cuticulome.org custom HMMs, an overall confidence level, and the supporting evidence (**Figure 2D**). Based on these results, a user can move from having an unannotated or poorly annotated arthropod protein sequence to having a more informed interpretation of its CP identity, family classification, related proteins, and even a possible function. While useful for all arthropod species, this could be particularly useful for data proceeding from non-model organisms that have incomplete sequence annotation or lack a well-defined genome, transcriptome, or proteome.

**Figure 2.**
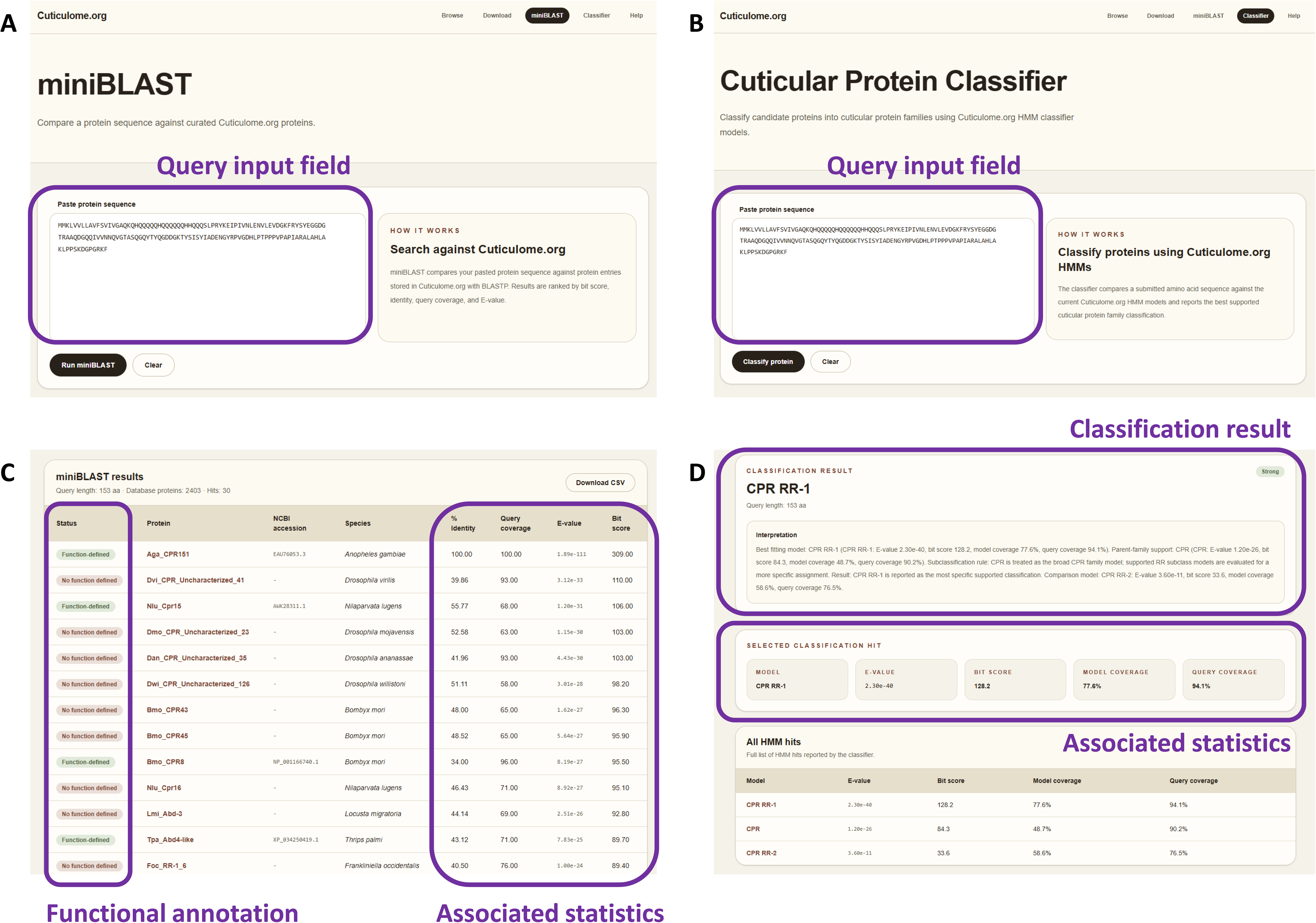
Characterization and classification of a putative CP use-case. (A) Query of CP sequence in raw format implementing miniBLAST. (B) Query of CP sequence in raw format in Cuticular Protein Classifier. (C) miniBLAST match results with additional related statistics, annotation information, and function-defined status indicator. (D) Cuticular Protein Classifier results including established CP family classification and associated HMM statistics. As an example, *A. gambiae* CPR151 was used in these queries.

### Example 2: “Plug and play” datasets

Researchers that study related phylogeny or carry out comparative analyses such as the evolution of a CP family, the expansion of CP repertoires in specific arthropod lineages, or the relationship between CPs of similar functions can use Cuticulome.org to have ready-to-use datasets within just a few clicks (**Figure 3**). This avoids troublesome collection of individual sequences that are scattered across independent publications, species-specific databases, and general protein databases that do not conserve nomenclature or functional annotations.

**Figure 3.**
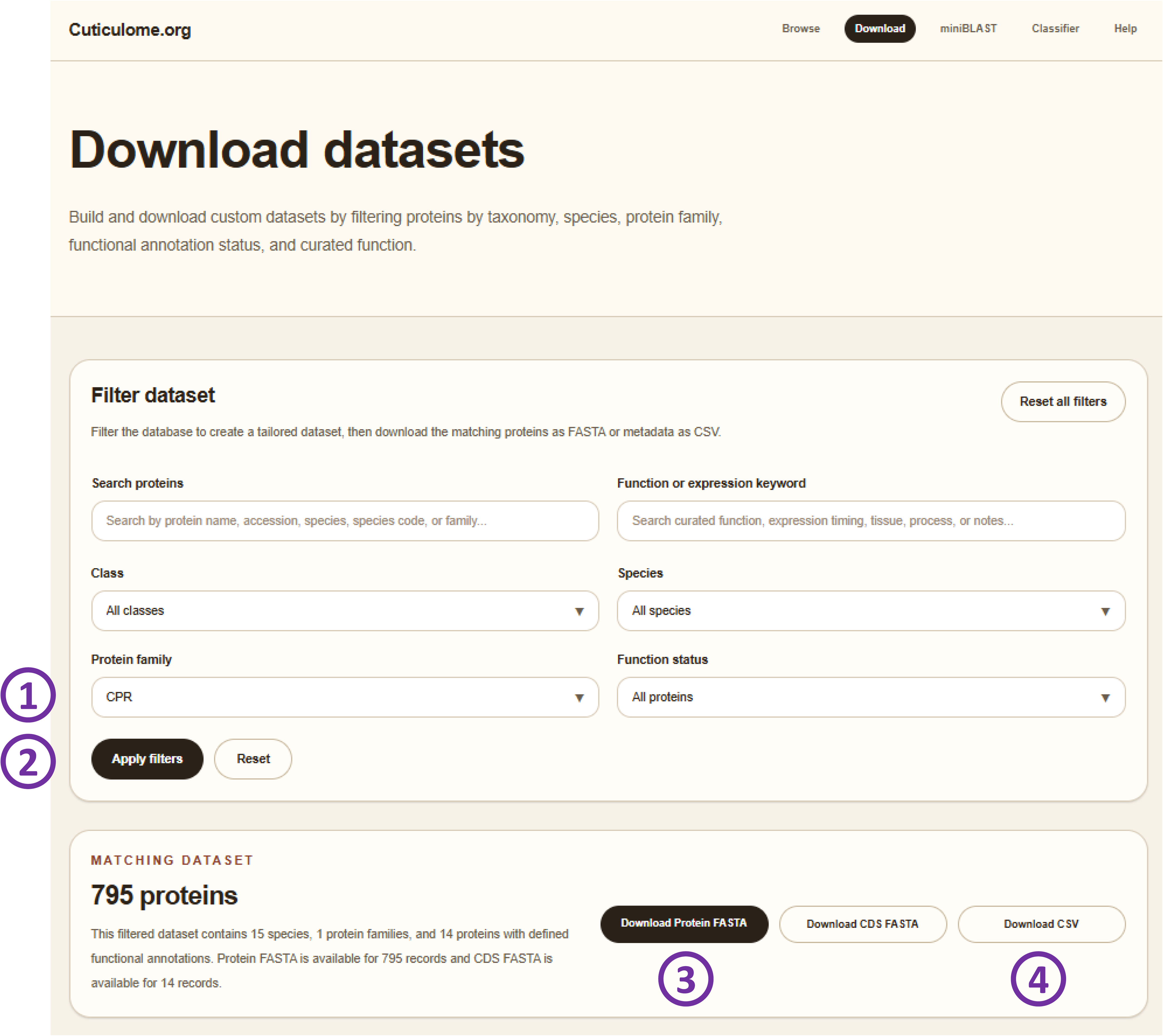
Building and downloading a customized dataset use-case. To download a specific protein family dataset, users would first select the established protein family of interest (1), apply the filters (2), and then download the filtered protein FASTA (3) and associated annotations in CSV format (4).

As an example, a researcher interested in investigating the evolution of a specific CP family can access the Download page and filter the database based on the protein family of interest **(Figure 3A**). After applying filters (**Figure 3B**) they can obtain all amino acids of a specific CP family with standardized headers in FASTA format that can be directly used in a phylogenetic analysis (**Figure 3C**). Following the results of a phylogenetic tree, they can investigate the function relation of a specific branch, which they can easily browse in the CSV they can download simultaneously from the filtered dataset (**Figure 3D**). In this way, Cuticulome.org allows users to fluidly move through datasets of interest without the need for difficult editing and re-editing of FASTA files or functional notes.

## FUTURE DIRECTIONS

Cuticulome.org offers a modular design that can accommodate future tools and data updates. In instance, submitting new data via the portal submission could be a quick approach to expand the database. We also envision to develop and incorporate modern AI-based structural predictions. Our database also allows us to visualize big evolutionary coverage gaps. In the coming years, we expect to see a significant rise in CP data and knowledge in underrepresented subphyla such as Chelicerata (spiders, scorpions), Crustacea (crabs, shrimps), and Myriapoda (centipedes, millipedes). Moreover, CPs hold the potential to transform the chitosan-based material landscape, which should grow the “Engineered CP” family.

## DATA AVAILABILITY

Cuticulome.org is freely accessible at https://www.Cuticulome.org. The source code, curated data tables, and web interface implementation are available from the dedicated Cuticulome.org GitHub repository at https://github.com/Wardale24/Cuticulome_web.

## ACKNOWLEDGEMENTS

We would like to thank Gonzalo Merino Pérez-Arroyo for thoroughly inspecting, testing, and suggesting valuable web development changes and additions to the Cuticulome.org website. We additionally thank all researchers that have generated and made protein data publicly available for the use by the international research community, as well as those who were in private communication allowing us to incorporate their datasets into Cuticulome.org.

## AUTHOR CONTRIBUTIONS

Conceptualization: A.W., C.F.; Data Curation: A.W., C.F.; Formal analysis: A.W.; Funding acquisition: N.N.; Investigation: A.W., C.F.; Methodology: A.W., C.F.; Software: A.W.; Supervision: C.F; Validation: A.W., C.F.; Visualization: A.W., C.F.; Writing — Original Draft: A.W., C.F.; Writing — Review & Editing: A.W., N.N., C.F.

